# The crystal structure of the human SIPA1L2 PDZ domain reveals an atypically short, metazoan-conserved β2–β3 loop

**DOI:** 10.64898/2026.11.16.766968

**Authors:** Torben Klos, Boglarka Zambo, Alastair G. McEwen, Bastien Morlet, Gergo Gogl, Helgo Schmidt

**Author notes:** Correspondence: Gergo Gogl,; Helgo Schmidt.

## Abstract

Signal-induced proliferation-associated 1-like protein 2 (SIPA1L2) is a large neuronal Rap GTPase-activating protein implicated in dynein-dependent retrograde transport of BDNF– TrkB signaling endosomes and genetically linked to Parkinson’s disease. It consists of a RapGAP domain, a PDZ domain, and a coiled-coil domain separated by intrinsically disordered regions. Its compact PDZ domain mediates protein-protein interaction by binding short PDZ-binding motifs (PBM) in target proteins. Here, the human SIPA1L2 PDZ domain (residues 943–1027) was crystallized and its structure determined by molecular replacement. The domain adopts the canonical PDZ fold of six β-strands forming two antiparallel β-sheets flanked by two α-helices. The β2–β3 loop, located next to the PBM binding groove, is unusually short — eight residues, against a 14-residue human-PDZome consensus. A BLASTP-and AlphaFold3-based survey across Metazoa shows that, despite PDZ sequence identities falling to ∼25 %, the fold and the short loop are structurally conserved (Cα RMSD 0.5–0.8 Å), indicating that this is a genuine, evolutionarily constrained architectural feature of SIPA1L2-type PDZ domains. PBM binding was not observed in co-crystallization trials. Native holdup assays against a neuronal proteome identified no specific partners for the SIPA1L2 PDZ domain. The structure retains the canonical carboxylate-binding loop and a class-I-type histidine, so partner scarcity is not due to loss of the PBM binding apparatus. Impaired PBM binding seems to result from the combination of the unusually short β2–β3 loop and an evolutionarily invariant His964 located at the β2 strand. The short loop might restrict the conformational dynamics of the β2 strand, thereby stabilizing His964 side-chain conformations that encroach on the PBM binding groove. The structure thus defines SIPA1L2 as a structurally canonical yet apparently orphan PDZ domain.

**Synopsis:** The first experimental structure of the human SIPA1L2 PDZ domain adopts the canonical PDZ fold and carries an atypically short β2–β3 loop that is structurally conserved across Metazoa. A conserved histidine residue partially occludes the groove for the PDZ-binding motif.

## 1. Introduction

Neurons depend on long-range retrograde transport to relay neurotrophic signals from distal axons to the soma. Brain-derived neurotrophic factor (BDNF), bound to its receptor TrkB, is internalized into signaling endosomes that are carried toward the soma by cytoplasmic dynein. The cargo-adaptor machinery that couples BDNF–TrkB carriers to dynein remains incompletely defined (Ha et al., 2008; Villarroel-Campos et al., 2018). Among candidate adaptors, the multidomain protein SIPA1L2 (signal-induced proliferation-associated 1-like protein 2), a member of the SIPA1 family of Rap GTPase-activating proteins, bridges TrkB to a dynein motor through the cargo adaptor Snapin (Andres-Alonso et al., 2019). It was demonstrated that Snapin bound to a SIPA1L2 fragment spanning the RapGAP and PDZ domains (residues 470–1025). The isolated PDZ domain has never been shown to bind Snapin or any other ligand. Its role was left open, with the suggestion that it might dock the complex to presynaptic sites through interactions between the PDZ domain and PDZ-binding motifs (PBM) (Andres-Alonso et al., 2019). SIPA1L2 is also of medical interest. The rs10797576 intronic SIPA1L2 variant is a genome-wide-significant risk locus for Parkinson’s disease (Nalls et al., 2014; Chang et al., 2017).

SIPA1L2 is a large and predominantly intrinsically disordered protein. Its N-terminal Rap-GAP domain, a single PDZ domain, and C-terminal coiled-coil α-helices are the only regions predicted to be well-folded. PDZ domains are compact (∼90-residue) protein–protein interaction modules that typically recognize short C-terminal peptide PBMs in a groove between strand β2 and helix α2 (Doyle et al., 1996; Morais Cabral et al., 1996; Ernst et al., 2014). Notably, high-throughput screens of the human PDZome have returned essentially no high-confidence PBM partners for the SIPA1L2 PDZ domain (Gogl et al., 2022; Vincentelli et al., 2015). The structural basis for this apparent absence of canonical binding partners is unknown. Here we report the first experimental structure of a SIPA1L2 PDZ domain in three crystal forms, describe its canonical fold featuring an unusually short β2– β3 loop, use a structure-guided phylogenetic survey to show that this short loop is conserved across Metazoa, and assess whether the apo groove geometry can account for the domain’s impaired PBM binding behavior.

## 2. Materials and methods

### 2.1. Macromolecule production

The human SIPA1L2 PDZ domain (UniProt Q9P2F8, residues 943–1027) was expressed in *Escherichia coli* C41(DE3) as a fusion carrying an N-terminal His₆ tag followed by a TEV-protease cleavage site and a C-terminal human annexin A2 (ANXA2) crystallization chaperone lacking its disordered N-terminus, following the ANXA2-assisted strategy of Cousido-Siah *et al*. (2022) to aid the crystallization of highly soluble PDZ domains (Table 1).

**Table 1.** Macromolecule-production information.

|  |  |
|---|---|
| Source organism | *Homo sapiens* (SIPA1L2 PDZ domain; ANXA2 chaperone) |
| DNA source | Synthetic gene, codon-optimized for insect-cell and bacterial expression (Epoch Life Science) |
| Expression vector | Modified pET-15b (T7 promoter): N-terminal His<sub>6</sub> tag and C-terminal ANXA2 (residues 23–339) crystallization chaperone; Addgene plasmid #136546 |
| Expression host | *E. coli* C41(DE3) |
| Complete amino-acid sequence of the construct produced | MGHHHHHHDYDIPTTENLYFQGSHMTRGCETVEMTLRRNGLGQLGFHVNFEGIVADVEPFGFAWKAGLRQGSRLVEICKVAVATLTHEQMIDLLRTSVTVKVVIIQPHDDGSAYGSVKAYTNFDAERDALNIETAIKTKGVDEVTIVNILTNRSNEQRQDIAFAYQRRTKKELASALKSALSGHLETVILGLLKTPAQYDASELKASMKGLGTDEDSLIEIICSRTNQELQEINRVYKEMYKTDLEKDIISDTSGDFRKLMVALAKGRRAEDGSVIDYELIDQDARDLYDAGVKRKGTDVPKWISIMTERSVPHLQKVFDRYKSYSPYDMLESIRKEVKGDLENAFLNLVQCIQNKPLYFADRLYDSMKGKGTRDKVLIRIMVSRSEVDMLKIRSEFKRKYGKSLYYYIQQDTKGDYQKALLYLCGGDD* |

Expression was induced at an OD600 of 0.5–0.8 with 1 mM IPTG, and the culture was grown overnight (≈16 h) at 18°C. Harvested cells were resuspended in 50 mM HEPES pH 7.5, 150 mM NaCl, 5% glycerol, 0.1% Triton X-100, 1 mM MgATP, 20 mM imidazole, 5 mM β-mercaptoethanol (BME), 2 mM PMSF, cOmplete EDTA-free protease-inhibitor cocktail (Roche) and a trace of DNase I and lysozyme; the resuspended cells were frozen at −20°C and thawed on ice before purification (or, when purification was performed the same day, frozen briefly at −80°C for 1 h). Cells were lysed by sonication (5 × 45 s pulses at 60% amplitude with 30 s pauses; duty cycle 0.5) and clarified by ultracentrifugation (148 000 × g, 30 min, 4°C). The clarified supernatant was applied to a pre-equilibrated Ni²⁺-charged HisTrap HP column (Cytiva), washed with buffer containing 30 mM imidazole and eluted with 400 mM imidazole (20 mM HEPES pH 7.5, 5% glycerol, 150 mM NaCl, 5 mM BME); the wash and elution buffers were held at slightly reduced pH to prepare the sample for subsequent cation-exchange chromatography. Protein-containing fractions were pooled and incubated with TEV protease for 1–2 h at room temperature to remove the His₆ tag, then loaded onto a HiTrap SP HP cation-exchange column and eluted with a linear NaCl gradient. Peak fractions were pooled, concentrated, and buffer-exchanged on a HiTrap Desalting column (Cytiva) into 20 mM HEPES pH 6.5, 150 mM NaCl, 2 mM CaCl₂, 2 mM TCEP, supplemented with 20 mM CaCl₂ for the Li and Ca forms (Table 2). Final samples were concentrated to 5–7 mg ml⁻¹, aliquoted, flash-frozen in liquid nitrogen, and stored at −80°C.

**Table 2.** Crystallization parameters.

| Method | Sitting-drop vapor diffusion |
|---|---|
| Plate type | 96-well 2-drop MRC (SWISSCI) |
| Temperature (K) | 293 |
| Protein concentration | 60 µM (fusion) |
| Ligand in the drop | Snapin-derived phosphopeptide Biotin-ttds-HVHAVRE(pSer)QVELREQ, 174 µM (2.9-fold molar excess over the fusion) |
| Buffer composition of protein solution | 20 mM HEPES pH 6.5, 150 mM NaCl, 2 mM TCEP (noCa form); the same buffer supplemented with 20 mM CaCl<sub>2</sub> (Li and Ca forms) |
| Composition of reservoir solution | 0.2 M tri-lithium citrate, 20 %(w/v) PEG 3350 |
| Volume and ratio of drop | 150 nL + 150 nL (1:1) |
| Volume of reservoir | 40 µL |

### 2.2. Crystallization

Our initial aim was to elucidate whether Snapin is able to bind to the SIPA1L2 PDZ domain. To this end, we carried out co-crystallization experiments with a Snapin-derived peptide (residues 43 - 57). The peptide is a 15-mer centered on the Ser50 PKA site of Snapin. This Snapin peptide region also interacts with the dynein intermediate chain. PKA phosphorylates Snapin at Ser50 (Chheda et al., 2001), and PKA activity releases the SIPA1L2–Snapin complex from the dynein intermediate chain (Andres-Alonso et al., 2019), which raised the possibility that Ser50-phosphorylated Snapin is the form available for interaction with the SIPA1L2 PDZ domain. We therefore focused our co-crystallization attempts on the Ser50-phospho peptide. Purified SIPA1L2 PDZ–ANXA2 fusion and the Snapin-derived phosphopeptide (Biotin-ttds-HVHAVRE(pSer)QVELREQ) were combined to final concentrations of 60 µM and 174 µM, respectively — a 2.9-fold molar excess of peptide — and the mixture was used directly for crystallization. Sitting-drop vapor-diffusion experiments were performed at 293 K using a Mosquito Crystal nanoliter dispenser (SPT Labtech) by mixing 150 nL protein with 150 nL reservoir solution and equilibrating against 40 µL reservoir in 96-well 2-drop MRC plates (SWISSCI). A total of 288 conditions were screened (commercial JCSG+ and PACT screens and an in-house TOP96 screen; Qiagen, Hampton Research, Molecular Dimensions and Rigaku Reagents). Crystals appeared within ∼1 h. Diffraction-quality crystals grew in 0.2 M tri-lithium citrate, 20 %(w/v) PEG 3350. Crystallization parameters are summarized in Table 2. Crystals were cryoprotected in reservoir supplemented with 35 %(w/v) PEG 3350 and flash-cooled in liquid nitrogen.

We obtained crystals under three different ion states that arose from two protein-buffer compositions equilibrated against the same tri-lithium citrate/PEG 3350 reservoir. The noCa form grew from protein in 20 mM HEPES pH 6.5, 150 mM NaCl, 2 mM TCEP. For the Li and the Ca form, the protein buffer was supplemented with 20 mM CaCl_2_. In the resulting P2₁2₁2₁ (Li) lattice, the chaperone sites were nevertheless occupied by Li⁺ from the 0.2 M tri-lithium citrate reservoir, whereas in the C2 (Ca) lattice the same sites were occupied by Ca²⁺ (Fig. S1, Section S1).

### 2.3. Data collection and processing

Diffraction data were collected from single crystals at 100 K. The Li and Ca forms were measured on beamline PROXIMA 1 (Synchrotron SOLEIL, Saint-Aubin, France) at a wavelength of 0.9786 Å using an EIGER-X 16M detector (Dectris), and the noCa form on beamline ID30B (ESRF, Grenoble, France) at 0.9116 Å using an EIGER2 × 9M detector (Dectris). Data were indexed, integrated, scaled, and merged with *autoPROC* (Vonrhein et al., 2011). The fusion protein crystallized in three ion states — referred to here as noCa, Li and Ca — diffracting to 1.61, 1.50 and 2.01 Å, respectively. The Li data are essentially isotropic and complete (99.94 %). The noCa and Ca data are strongly anisotropic and were therefore corrected by anisotropic truncation and Bayesian scaling in *STARANISO* (Tickle et al., 2018), which accounts for the apparent deficit in their spherical completeness (74.6 % and 61.0 %). Along the best-diffracting direction, the noCa data extend to 1.61 Å and the Ca data to 2.01 Å, while the worst directions are limited to 2.03 and 2.92 Å. The corresponding ellipsoidal completeness is 94.1 % and 90.7 %. Refinement was stable for all three forms, and CC₁⁄₂ remains high over most of the resolution range (Table 3). All three forms are reported to document the ion-dependent lattice variation; the high-resolution Li form serves as the reference model throughout. Statistics are given in Table 3.

**Table 3.** Data collection and refinement statistics Values in parentheses are for the outer (highest-resolution) shell.

|  | noCa (P2<sub>1</sub>2<sub>1</sub>2<sub>1</sub>) | Li (P2<sub>1</sub>2<sub>1</sub>2<sub>1</sub>) | Ca (C2) |
|---|---|---|---|
| **Data collection** |  |  |  |
| Beamline | ID30B (ESRF) | PROXIMA-1 (SOLEIL) | PROXIMA-1 (SOLEIL) |
| Detector | EIGER2 X 9M (Dectris) | EIGER-X 16M (Dectris) | EIGER-X 16M (Dectris) |
| Wavelength (Å) | 0.9116 | 0.9786 | 0.9786 |
| Space group | P2<sub>1</sub>2<sub>1</sub>2<sub>1</sub> | P2<sub>1</sub>2<sub>1</sub>2<sub>1</sub> | C2 (C121) |
| *a, b, c* (Å) | 42.59, 93.94, 124.53 | 42.73, 93.88, 124.79 | 130.997, 61.882, 60.619 |
| *α, β, γ* (°) | 90, 90, 90 | 90, 90, 90 | 90, 97.354, 90 |
| Resolution (Å) | 51.9–1.61 (1.668–1.61) | 30.63–1.50 (1.554–1.50) | 64.96–2.012 (2.084–2.012) |
| Total reflections | 1398919 (106775) | 872904 (87552) | 210210 (20339) |
| Unique reflections | 125062 (996) | 155255 (8002) | 62050 (73) |
| Multiplicity | 11.2 (8.5) | 5.6 (5.6) | 3.4 (3.3) |
| Completeness (%) | 74.60 (15.38) | 99.94 (99.84) | 60.96 (2.30) |
| ⟨I/σ(I)⟩ | 7.05 (0.34) | 10.96 (0.80) | 3.11 (0.30) |
| Wilson B (Å<sup>2</sup>) | 19.42 | 20.23 | 26.27 |
| R_merge | 0.536 (8.350) | 0.086 (1.896) | 0.497 (5.144) |
| R_meas | 0.562 (8.885) | 0.095 (2.091) | 0.592 (6.151) |
| R_pim | 0.169 (3.006) | 0.040 (0.874) | 0.318 (3.343) |
| CC<sub>1</sub>/<sub>2</sub> | 0.984 (0.075) | 0.999 (0.418) | 0.945 (0.002) |
| CC* | 0.996 (0.375) | 1.000 (0.768) | 0.986 (0.056) |
| Anisotropic truncation |  |  |  |
| Ellipsoidal resolution (Å) | 2.03 (a*), 1.61 (b*), 1.63 (c*) | n/a | 2.92 (0.989a* + 0.148c*), 2.01 (b*), 2.06 (−0.683a* + 0.730c*) |
| Unique reflections (ellipsoidal) | 49008 (2451) | n/a | 19577 (741) |
| ⟨I/σ(I)⟩ (ellipsoidal) | 13.0 (1.5) | n/a | 6.8 (1.4) |
| R_meas (ellipsoidal) | 0.386 (8.941) | n/a | 0.335 (1.630) |
| R_pim (ellipsoidal) | 0.082 (1.982) | n/a | 0.129 (0.633) |
| CC<sub>1</sub>/<sub>2</sub> (ellipsoidal) | 0.999 (0.692) | n/a | 0.984 (0.359) |
| Ellipsoidal completeness (%) | 94.1 (69.7) | n/a | 90.7 (46.5) |
| **Refinement** |  |  |  |
| Reflections, working set | 49003 (996) | 81246 (7996) | 19577 (73) |
| Reflections, test set (R_free) | 2394 (61) | 4061 (400) | 1007 (5) |
| R_work | 0.168 (0.321) | 0.125 (0.304) | 0.189 (0.221) |
| R_free | 0.198 (0.404) | 0.169 (0.332) | 0.240 (0.353) |
| CC(work) | 0.943 (0.823) | 0.977 (0.758) | 0.933 (0.778) |
| CC(free) | 0.942 (0.764) | 0.961 (0.671) | 0.940 (0.828) |
| No. non-H atoms | 3833 | 4230 | 3507 |
| protein | 3250 | 3423 | 3231 |
| ligand/ion | 24 | 50 | 96 |
| solvent | 559 | 757 | 180 |
| Protein residues | 400 | 401 | 400 |
| R.m.s.d. bonds (Å) | 0.003 | 0.005 | 0.005 |
| R.m.s.d. angles (°) | 0.52 | 0.81 | 0.66 |
| Ramachandran favored (%) | 98.24 | 98.50 | 96.98 |
| Ramachandran allowed (%) | 1.26 | 1.25 | 2.27 |
| Ramachandran outliers (%) | 0.50 | 0.25 | 0.76 |
| Rotamer outliers (%) | 0.28 | 0.26 | 0.84 |
| Clashscore | 1.66 | 2.66 | 4.29 |
| Average B (Å<sup>2</sup>) | 24.64 | 26.97 | 34.22 |
| TLS groups | 4 | none (anisotropic ADPs) | 3 |
| Matthews coeff. (Å<sup>3</sup> Da<sup>−1</sup>) / solvent (%) | 2.69 / 54.3 | 2.71 / 54.5 | 2.63 / 53.3 |
| PDB code | 35IQ | 35IS | 35IT |

### 2.4. Structure solution and refinement

The noCa structure was solved by molecular replacement with *Phaser* (McCoy et al., 2007) using a two-body search: an AlphaFold-predicted model of the SIPA1L2 PDZ domain (Jumper et al., 2021; Varadi et al., 2024) and the ANXA2 core taken from PDB entry 7p73 (Gogl et al., 2022). The Li form is isomorphous with the noCa form, and its structure was obtained by direct refinement of the noCa model against the lithium data rather than by a separate molecular-replacement search. The Ca form, which crystallized in a different space group, was solved by molecular replacement using the refined PDZ and ANXA2 components of the Li structure as independent search models. Models were rebuilt in *Coot* (Emsley et al., 2010) and refined in *phenix.refine* (Adams et al., 2010) through iterative cycles of manual rebuilding and restrained refinement. TLS parameterization was used for the noCa and Ca forms; the Li form was refined without TLS, with anisotropic displacement parameters for all non-hydrogen atoms, as permitted by its resolution and completeness. Waters, ligands, and ions were placed from F_o−F_c and 2F_o−F_c difference maps and assessed by coordination geometry and refinement behavior. Metal-site coordination spheres and bond-valence sums were calculated, including crystallographic symmetry mates, using bond-valence parameters (Brown, 2009), and cross-validated with *CheckMyMetal* (Zheng et al., 2017), which evaluates the deposited asymmetric unit. Model quality was assessed by R_work/R_free and MolProbity (Williams et al., 2018) to generate Ramachandran, rotamer, and clashscore statistics. Figures were prepared with PyMOL (Schrödinger LLC, 2015).

### 2.5. Sequence and structural analyses

β2–β3 loop boundaries were assigned on a multiple sequence alignment of human PDZ domains rather than from the secondary structure of individual models, so that loop length could be compared across the PDZome, including domains without experimental structures. The alignment contains 258 of the 266 human PDZ domains assayed in ProfAff (Gogl et al., 2022). The four HtrA PDZ domains, which lack the canonical carboxylate-binding loop that anchors the loop definition, and four domains whose alignment in the loop region was ambiguous (CNKSR1, INTU, PDZD2 PDZ1 and DEPDC2 PDZ2) were excluded. Under this convention, the SIPA1L2 loop comprises eight residues (Phe963– Ile970, FHVNFEGI); DSSP (Kabsch & Sander, 1983) assigns Phe963–Val965 to β2 and Val971–Val974 to β3, so the structurally defined loop is Asn966–Ile970, with Ile970 forming an isolated β-bridge. PDZ:PBM affinities were taken from the ProfAff dataset (Gogl et al., 2022). A domain was counted as a binder if at least one interaction was quantified with pKd > 4.5 (Kd < 32 μM), a threshold above the quantification limit of all individual holdup assays. Because 129 of the 258 domains were screened against the full panel of 448 PBMs and the remainder against a subset of 40–45 PBMs, the correlation analysis was restricted to the 129 fully screened domains (point-biserial correlation and logistic regression for binary PBM competence; Pearson and Spearman correlation for partner counts among domains with at least one partner). Short-looped PDZ domains were identified from the alignment and, where structures were available, superposed on the experimental SIPA1L2 structure in UCSF ChimeraX (Meng et al., 2023). Comparators with available structures were APBA3 PDZ2 (Dang et al., 2008) and SDCBP/syntenin PDZ2 (Bradshaw et al., 2022). For the metazoan survey, representative SIPA1L2 homologs were retrieved by BLASTP against UniProtKB Reference Proteomes and Swiss-Prot (BLOSUM62, *E* ≤ 10, ≤250 alignments, low-complexity filtering disabled). Per-taxon PDZ regions were modeled with AlphaFold3 (Abramson et al., 2024) and superposed (Cα, ChimeraX matchmaker) to obtain backbone RMSD.

Apo-groove geometry was analyzed from the refined coordinates of all three forms. The PDZ domains (residues 950–1024) were superposed onto the Li form (Cα; Biopython; Cock et al., 2009), and per-residue Cα and side-chain atom deviations were computed, providing a within-lattice (noCa) and across-lattice (Ca/C2) comparison. To test peptide-binding compatibility, five PDZ:PBM complexes (PDB 4e3b (Amacher et al., 2013), 4k72 (Amacher et al., 2014), 5ovc (Ponna et al., 2018), 7qct (Zhu et al., 2022), 2kpl (Charbonnier et al., 2011) were each superposed onto the SIPA1L2 PDZ domain of each crystal form by TM-align (Zhang & Skolnick, 2005), the resulting transformation applied to the bound peptide, and atom clashes (<2.5 Å) between the transferred peptide and SIPA1L2 enumerated and partitioned into main-chain and side-chain contributions (SciPy cKDTree; Virtanen et al., 2020). Disordered atoms were represented by conformer A. Peptide main-chain atoms within 2.5 Å of SIPA1L2 main-chain atoms were counted as main-chain clashes, except contacts between the peptide C-terminal carboxylate oxygens and the backbone amide N atoms of Leu961–Phe963, which correspond to the anchoring hydrogen bonds and are reported separately. Correct docking was confirmed by the proximity of the peptide C-terminus to the SIPA1L2 carboxylate-binding loop. Induced-fit movement in a reference binder was measured by superimposing apo (5ova; Ponna et al., 2018) and peptide-bound (5ovc; Ponna et al., 2018) Shank3 PDZ domains. The β2–β3-loop residue facing the peptide channel was identified by transferring the docked iCAL36-L peptide (from 4e3b) and measuring atom-level contacts. The structurally equivalent residue in binders was taken as the nearest Cα after TM-alignment onto SIPA1L2.

### 2.6. Native holdup assay

Partner identification used the native holdup (nHU) assay (Zambo et al., 2022). The bait was AviTag-MBP fused via a flexible deca-asparagine linker to the SIPA1L2 PDZ domain with a C-terminal His₆ tag. The control was biotinylated AviTag-His_₆_-MBP alone. Baits and controls were co-expressed with BirA in *E. coli* BL21 and were purified by two-step Ni-NTA and amylose affinity chromatography. The PDZ boundaries were identical (residues 943– 1027) in the nHU bait and in the crystallized construct. The bait or control was biotinylated (BirA), immobilized on streptavidin–Sepharose, and incubated (≈2 h, 4 °C) with a cleared SH-SY5Y neuroblastoma extract prepared in EDTA-free buffer (50 mM HEPES-KOH pH 7.5, 150 mM NaCl, 1 % Triton X-100, 5 mM TCEP, 10 % glycerol, Roche Protease inhibitor cocktail; EDTA omitted to preserve metal-ion-mediated interactions). Unbound prey in the supernatant was quantified by nanoLC-MS/MS in technical triplicate (n = 3 bait versus n = 3 control). Per-protein depletion (log₂ fold-change versus control) and significance (two-tailed unpaired Student’s *t*-test) were combined with a dataset-adaptive hyperbolic threshold (minimum −log₁₀P = 1.3; minimum fold-change = 1σ, with a curvature parameter of 1 as described in Zambo et al., 2022) to classify interactors, with apparent affinities (p*K*_app_) estimated as in Keilhauer et al., 2015. Gene Ontology / functional enrichment of the hit list was assessed with GOrilla (Eden et al., 2009) and DAVID (Sherman et al., 2022). Quantitative affinity comparisons across the SIPA1/SIPA1L family (SIPA1 Q96FS4 fragment 683–764, SIPA1L1 O43166 fragment 946–1030, SIPA1L2 Q9P2F8 fragment 943–1027, SIPA1L3 O60292 fragment 959–1043) were taken from the ProfAff database(Gogl et al., 2022). PBM interactions were judged by combined significance (−log10P) and binding-detection ratio in addition to p*K*_app_.

## 3. Results and discussion

### 3.1. Overall structure

The SIPA1L2 PDZ domain was determined in three crystal forms from a single crystallization condition, distinguished by the ion content at the ANXA2 chaperone(Cousido-Siah et al., 2022): an ion-free form (noCa, P2₁2₁2₁, 1.61 Å), a lithium-bound form (Li, P2₁2₁2₁, 1.50 Å) (Fig. S1) and a calcium-bound form (Ca, C2, 2.01 Å) (Fig. S1). The two P2₁2₁2₁ forms have near-identical cells (*a* ≈ 42.6, *b* ≈ 93.9, *c* ≈ 124.6 Å). In contrast, the Ca form crystallized in C2 with a distinct cell, indicating that ion identity at the chaperone modulates lattice packing. A detailed analysis of the detected metal ion binding sites in the ANXA2 chaperone is provided in supplementary information (Section S1). Each asymmetric unit contains one PDZ–ANXA2 fusion. All models show excellent stereochemistry (Table 3). The Li form is essentially complete at 1.50 Å and provides the reference model used throughout.

The SIPA1L2 PDZ domain adopts the canonical PDZ fold: six β-strands (β1–β6) arranged into two antiparallel β-sheets flanked by two α-helices (α1, α2) and the canonical PBM peptide-binding groove between β2 and α2 (Fig. 1*a* and *b*). The defining structural feature of the SIPA1L2 PDZ domain is its short β2–β3 loop. With eight residues (FHVNFEGI in human SIPA1L2), it is roughly half the length of the 14-residue consensus loop derived from the human PDZome (FSLRGGIEPDEGIF) and sits at the short tail of a loop-length distribution centered on 14 residues across 258 human PDZ domains (Fig. S2). The experimental fold agrees closely with the AlphaFold2 prediction (Cα RMSD < 0.7 Å). The Snapin-derived phosphopeptide included in the crystallization setup (see Materials and Methods, 2.2 crystallization) was not observed in the maps of any form.

**Figure 1.**
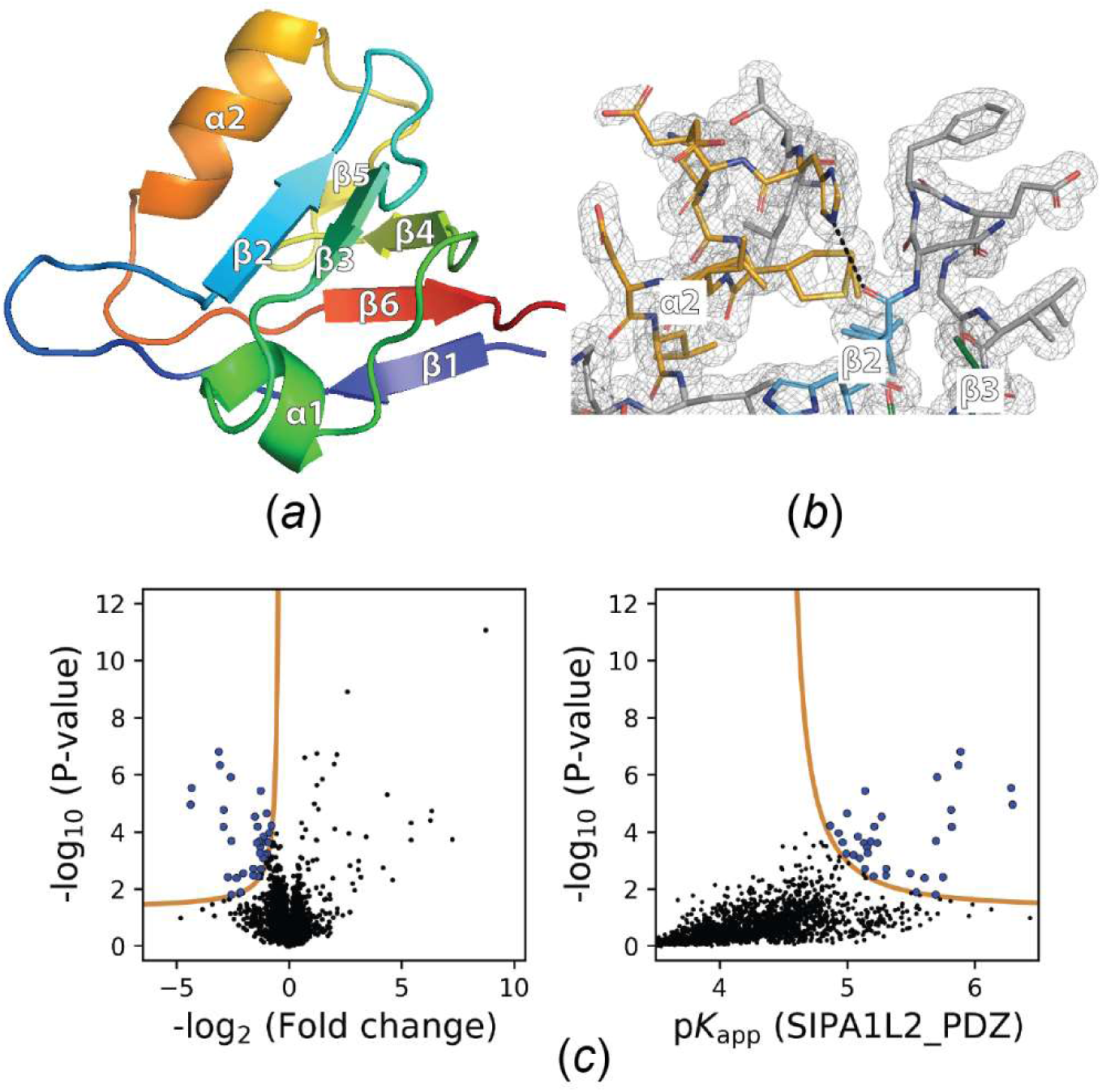
The SIPA1L2 PDZ domain features a binding groove impaired in PBM binding. (*a*) Overall structure of the SIPA1L2 PDZ domain (Li crystal form, 1.50 Å). Cartoon colored N→C with β1–β6 and α1–α2 annotated. The PBM binding groove lies between α2 and β2. It is empty despite the presence of a Snapin PBM in the crystallization buffer (see Materials and Methods, 2.2. Crystallization). **(*b*)** Example of the quality of the 2mFo−DFc electron density map contoured at 1σ. **(*c*)** Volcano plot of native holdup assay of the SIPA1L2 PDZ domain against a SH-SY5Y neuroblastoma cell lysate. Hits passing the significance threshold are colored in blue. However, none of the potential hits passed the additional validation criteria (see Section 3.2, main text), suggesting the SIPA1L2 PDZ domain is impaired in its PBM binding ability.

### 3.2. Native holdup assays and structural investigations suggest the SIPA1L2 PDZ domain features a partially blocked PBM binding groove

In order to elucidate the potential of the SIPA1L2 PDZ domain for PBM interactions in a complementary approach, we carried out a native holdup assay against a neuronal proteome (SH-SY5Y neuroblastoma cell line). In this assay, only 33 of 3,709 detected proteins exceeded the significance threshold (see Material and Methods, 2.6. Native holdup assay) (Fig. 1*c*). These hits were weak, showed no Gene Ontology or domain-architecture enrichment (GOrilla, DAVID), and included no recognizable C-terminal PBM partner, with the exception of ARHGAP17. In a previously reported complementary holdup screen against a defined PBM/PDZome panel, the SIPA1L2 PDZ domain bound no peptide with appreciable affinity(Gogl et al., 2022). This screen also included the isolated PBM of ARHGAP17, which showed no binding to SIPA1L2. We conclude that under the conditions tested and consistent with our co-crystallization results, the SIPA1L2 PDZ domain has no identified specific partner. Therefore, we examined the SIPA1L2 PDZ domain structure to investigate the molecular basis for this PBM binding deficiency.

When PBMs (PDB: 4e3b, 4k72, 5ovc, 7qct, 2kpl) are placed in the SIPA1L2 PBM binding groove of the Li crystal form by superposition (TM-scores 0.76–0.88) (Fig. 2*a*), the free C-terminal carboxylates sit 2.5–3.5 Å from the carboxylate-binding loop backbone amides (Leu961, Gly962, and Phe963), close to canonical carboxylate-anchoring distances. SIPA1L2 also carries a histidine (His1004) at the α2 position characteristic of class-I PDZ domains. In the docked structures, its imidazole ring lies 3.9 – 4.5 Å from the modeled P−2 residue of the four canonical peptides (PDB: 4e3b, 4k72, 5ovc, 7qct), not engaged but compatible with class-I P−2 contacts after induced fit. We obtain similar results when carrying out the PBM docking in the Ca-crystal form (Fig. S3). We observe no main-chain to main-chain clashes for the canonical complexes and five (Li form) and two (Ca form) clashes for the divergent MAGI-1/E6 complex (PDB: 2kpl) when comparing PBM docking in the Li-and Ca-crystal forms (Fig. 2*b*). Apart from His964 (see below), the residual peptide contacts (Fig. 2*b*) are minor side-chain clashes that could be relieved by induced fit. A bona-fide binder like Shank3 for example, reorganizes its PBM binding groove by up to 1.2 Å Cα shifts between apo (PDB: 5ova) and PBM bound (PDB: 5ovc) states, which is similar to what we observe between the SIPA1L2 PDZ P2₁2₁2₁ and C2 crystal forms (Fig. 2*c*). We conclude that the PBM binding groove of the SIPA1L2 PDZ domain has a carboxylate-binding loop in a suitable conformation as well as a class-I-type P−2 histidine to bind PBMs. The groove should also have, in general, sufficient plasticity to accommodate a PBM peptide. The lack of PBM binding partners for the SIPA1L2 PDZ domain can therefore not be explained by the absence of these PBM binding groove features.

**Figure 2.**
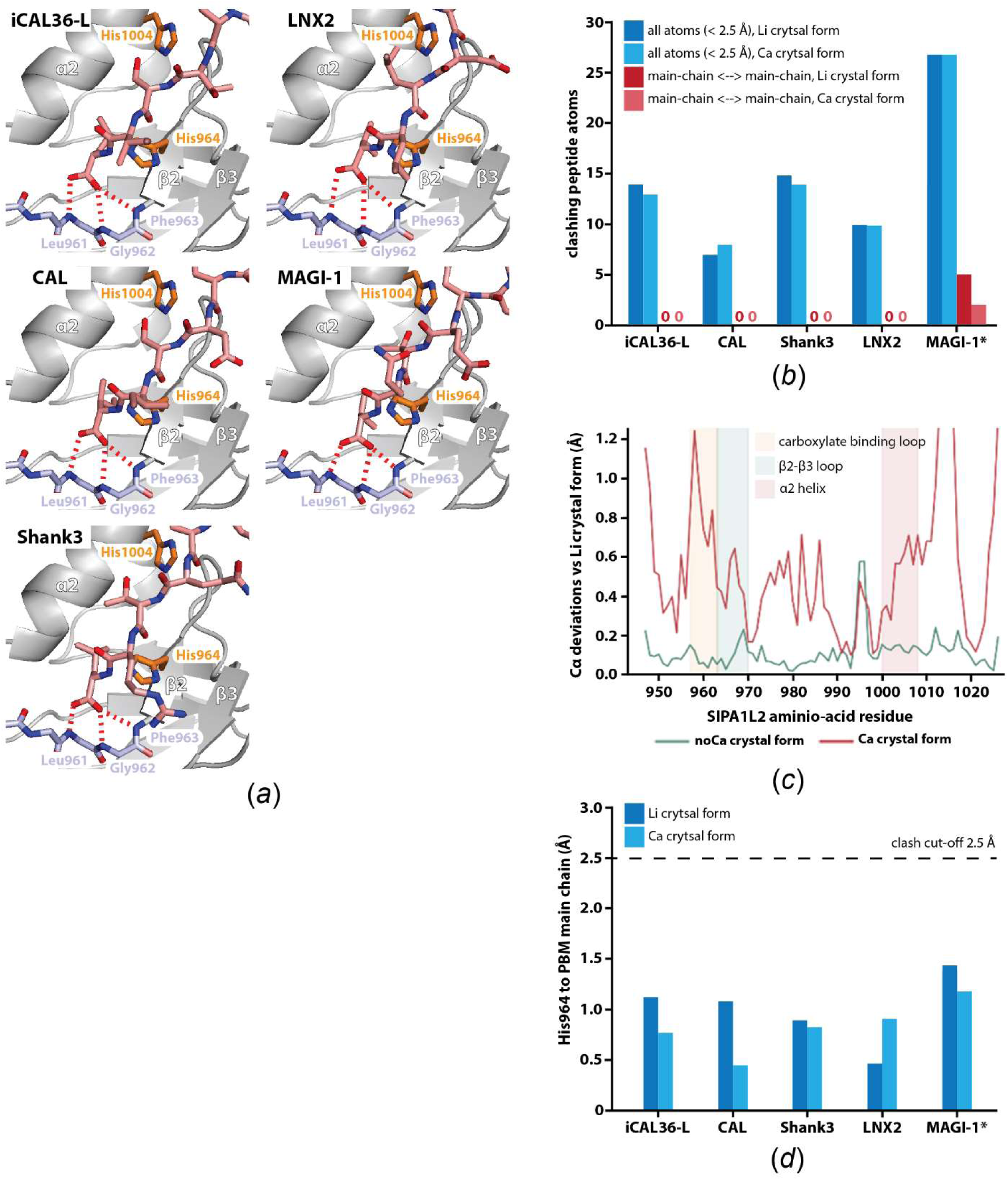
His964 in the SIPA1L2 PDZ domain partially obstructs the PBM binding groove. (*a*) To gain insight into SIPA1L2 PBM binding, we superimposed the PBMs (salmon) of iCAL36-L (PDB-ID: 4e3b), CAL (PDB-ID: 4k72), Shank3 (PDB-ID: 5ovc), LNX2 (PDB-ID: 7qct) and MAGI-1 (PDB-ID: 2kpl) onto the SIPA1L2 PBM binding groove. The SIPA1L2 PDZ domain (Li form) is shown in grey cartoon representation. The main chain of the carboxylate-binding loop (Leu961, Gly962, Phe963) is shown as light blue sticks. The α2 class-I reader His1004 and His964 are shown as orange sticks. The α2 helix and the β2 and β3 strands are labelled. In all cases, contacts with the canonical PBM binding motifs (carboxylate binding loop, α2 class-I reader) would be possible. Potential hydrogen bonds are shown as dashed red lines. A structural element that potentially would interfere with PBM binding is His964, which overlaps with the PBM main chain in all cases. **(*b*)** Steric clashes of the docked PBMs in the Li and Ca crystal forms (see also Fig. S3). **(*c*)** Structural plasticity of the SIPA1L2 PDZ PBM binding groove. Pairwise per-residue Cα deviation from the Li form to the same-lattice noCa form (P2₁2₁2₁, green) and the different-lattice Ca crystal form (C2, red). Structural elements surrounding the PBM binding groove are highlighted in light yellow (carboxylate binding loop), light green (β2-β3 loop), and light red (α2-helix). The P2₁2₁2₁ pair is essentially identical (∼0.1 Å), whereas the C2 form departs by up to ∼1.2 Å in the PBM groove area. Reduced structural plasticity is therefore unlikely to contribute to impaired PBM binding. **(*d*)** Distance between the His964 side chain and the PBM main chain in the docked PBM Li and Ca crystal form structures. In all docked complexes, His964 clashes with the PBM main chain.

The only obvious candidate interfering with PBM binding is His964, the second residue of strand β2. It projects towards the PBM binding groove and its imidazole side chain overlaps the modeled PBM peptide main chain in the Li-and Ca-crystal forms despite different His964 rotamers being present (Li: χ₁ −60°; Ca: χ₁ −174°) (Figs. 2*a* and S3). Across all docked PBM complexes, the His964 side chain approaches the peptide main chain to within 0.4–1.5 Å (Fig. 2*d*). In the canonical complexes no other groove residue comes closer than ∼2 Å. Only in the case of the divergent MAGI-1 complex another residue, Phe967, is closer with 0.5 – 0.7 Å. His964 is specific to the SIPA1L subfamily. In a curated alignment of the short-β2–β3-loop human PDZ domains, it is present at equivalent β2 positions only in SIPA1L1, SIPA1L2, and SIPA1L3. The PBM-binding short-loop PDZ domains have a small residue (Ser/Cis in the Mint/APBA domains, Ile/Val in the syntenins; Fig. S4), and across the whole human PDZome, a histidine is found in only 4.3 % (11/258) of domains. His964 is invariant in every sampled SIPA1L2 ortholog. Strikingly, this subfamily signature coincides with binding behavior. In a quantitative holdup screen between the human PDZ domains and 448 PBM peptides (ProfAff; Gogl et al., 2022), none of the three SIPA1L-family PDZ domains has a widespread PBM interactome. For SIPA1L2, only a handful of interaction partners were detected, all displaying ambiguous and marginal affinities. Out of the ∼25 detected interactions, only 5 displayed dissociation constants < 100 µM, whereas promiscuous PDZ domains normally capture several dozens to hundreds of partners with dissociation constants stronger than 10 µM in the same screen. The SIPA1L1 and SIPA1L3 PDZ domains were screened against smaller subsets of 40–43 PBMs and likewise yielded no partner above pKd 4.5. All three His964-bearing PDZ domains are thus orphans, whereas small-residue short-loop PDZ domains such as the syntenins, which do not have a His at the equivalent position, bind PBM peptides (Kang et al., 2003). However, this deficit is not exclusively confined to the His964-bearing branch. The more divergent paralog SIPA1, whose PDZ carries a glutamate at the His964 equivalent position, likewise yielded no partner in ProfAff (screened against 42 PBMs, Gogl et al., 2022). The absence of canonical PBM binding may therefore be a property of the whole SIPA1/SIPA1L PDZ clade and cannot be attributed to His964 alone. However, even though the identity of the side chain is not conserved, it is conceivable that the glutamate in SIPA1 sterically impairs PBM binding similar to His964 in SIPA1L2. Within the SIPA1L subfamily, His964 is the single conserved, PBM-binding groove-facing substitution that distinguishes these domains from the PBM-binding, short β2-β3 loop PDZs. His964 is therefore a structurally motivated candidate contributor to the PBM binding deficit in the SIPA1L subfamily.

### 3.3. The short loop β2–β3 loop is structurally conserved across Metazoa

To test conservation of the short β2–β3 loop, representative SIPA1L2 homologs were retrieved by BLASTP against UniProtKB reference proteomes and Swiss-Prot (BLOSUM62, *E* ≤ 10), the PDZ regions extracted, and AlphaFold3 models (Abramson et al., 2024) generated. The obtained models were superposed onto the experimental human SIPA1L2 PDZ domain structure (ChimeraX matchmaker, Cα). Although PDZ sequence identity declines steeply with phylogenetic distance — from 100 % in chimpanzee/mouse to ∼25 % in *Trichoplax* — backbone deviations remain small throughout (Cα RMSD 0.5–0.8 Å), and the eight-residue β2–β3 loop is retained across the sampled taxa (Table S1; Figs. 3*a* and 3*b*). This decoupling of structural from sequence conservation argues that the short β2–β3 loop is an evolutionarily constrained architectural feature of SIPA1L2-type PDZ domains. Sequence conservation within the loop is strongly position-dependent (Figs. 3*b* and 3*c*). Three positions are essentially invariant across SIPA1L2 orthologs — Phe963, His964, and Gly969. In contrast, the intervening positions are variable. The invariance of His964 is in particular interesting given its projection into the PBM binding groove.

**Figure 3.**
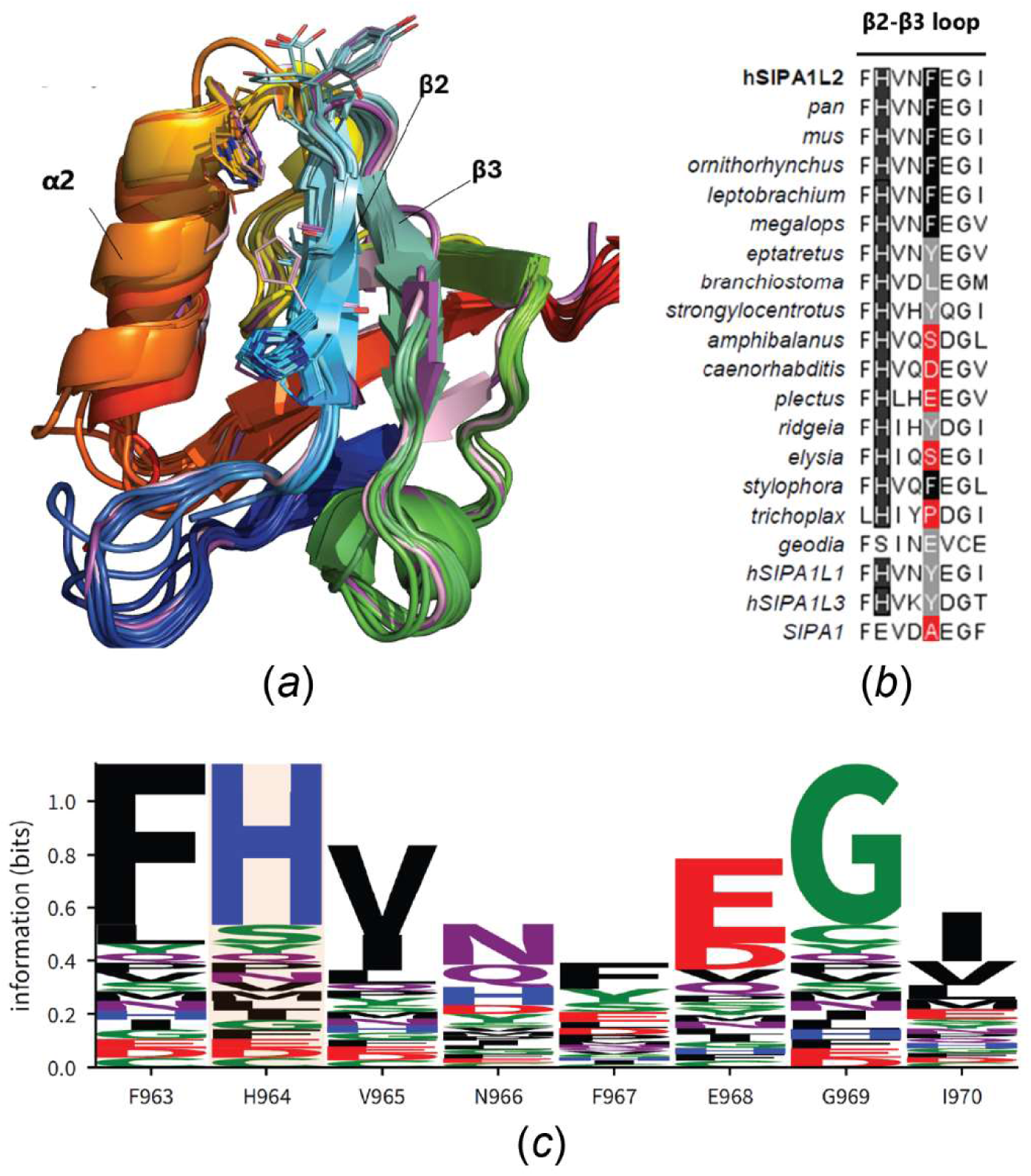
Conservation of the β2–β3 loop’s structure, length, and sequence. (*a*) Superposition of AlphaFold3 PDZ models (colored N→C) across taxa onto the experimental Li crystal form (purple) and Ca crystal form (pink) structures. **(*b*)** β2–β3-loop sequence alignment with conserved positions highlighted. **(*c*)** Sequence logo of the eight-residue β2– β3 loop across 17 metazoan homologs: Phe963, His964, and Gly969 dominate (invariant in SIPA1L2 orthologs).

### 3.4. Conclusions

To the best of our knowledge, we present here the first experimental structure of any SIPA1L2 domain. It provides a reference scaffold for interpreting SIPA1L2 sequence variation, including disease-associated variants, and a structural starting point for defining the PDZ domain ligand-recognition properties.

Our interactomics data revealed no obvious PBM binding partner for the SIPA1L2 PDZ domain, although it features all structural elements required for PBM binding. Also, our co-crystallization attempts with a Snapin PBM motif did not result in an occupied PBM binding groove. It is possible that the SIPA1L2 PDZ domain is orphan in the sense that suitable PBMs simply have not been identified yet.

There is also the possibility that the SIPA1L2 PDZ features a binding groove impaired in PBM interaction. It is quite striking that all SIPA1/SIPA1L PDZ clade family members investigated so far did not show PBM binding. Our combined structural analysis suggests that the conserved His964 contributes to the sealed PBM binding groove by sterically blocking access to it. However, since a His at a position equivalent to 964 is compatible with PBM binding in other PDZ domains, e.g., NHERF1 (Bhattacharya et al., 2013), it cannot be the only feature preventing PBM binding. In the Li-and noCa crystal forms, His964 is held in its groove-facing rotamer by a hydrogen bond to Asp973 (2.7–2.9 Å). In the Ca-crystal form, His964 adopts a different rotamer and the contact is broken.

The unusually short β2–β3 loop next to the PBM binding groove is another defining structural element of the SIPA1L2 PDZ domain. To investigate whether a short β2–β3 loop in general contributes to impaired PBM binding, we compared β2–β3 loop length with PBM binding across the 129 human PDZ domains screened against the complete ProfAff panel. Loop length does not predict whether a domain has a partner (point-biserial r = 0.12, p = 0.18; logistic regression odds ratio 1.08 per residue, 95 % CI 0.97–1.20). Among the 98 domains with at least one partner, longer loops are associated with somewhat more partners (Spearman ρ = 0.32, p = 0.0015), but the relationship is weak (R² = 0.06; Fig. S5). A short β2–β3 loop alone is therefore not sufficient to exclude PBM peptides from the binding groove. Other short-looped PDZ domains such as the syntenin PDZ2 bind PBM peptides well (Kang et al., 2003). However, short β2–β3 loops might nevertheless contribute to impaired PBM binding under more specific circumstances. For example, the exceptionally short SIPA1L2 PDZ β2–β3 loop might restrict dynamic movements of the β2 strand, which harbours His964, and therefore reinforce the steric exclusion of a PBM peptide.

Inhibition of PBM binding is a well-described phenomenon in the PDZ field. Prominent examples include proteins featuring a PDZ domain and a C-terminal PBM within the same protein. The C-terminal PBM then binds to the PBM-binding groove at the PDZ domain to block access. This increases the specificity of the PDZ domain, because only PBMs with a binding affinity higher than the binding affinity of the autoinhibited PBM can bind (Morales et al., 2007; Long et al., 2005). Also, steric occlusion of the PBM binding groove by neighbouring bulky side chains has been described. In the case of the Pals1 PDZ domain, a Phe residue partially obstructs the PBM binding groove, and the Ala substitution leads to a five-fold increase in PBM binding affinity (Ivanova et al., 2015). A related mechanism operates in the PTP1E PDZ2b splice variant, in which an extended β2–β3 loop folds back onto the binding site (Kachel et al., 2003).

What could be the biological purpose of an inhibited SIPA1L2 PDZ domain? An obvious benefit of such an inhibition would be the precise timing of protein-protein interactions upon signals triggering conformational changes in the PDZ domain to overcome PBM binding groove blockage. The SIPA1L2-mediated dynein-dependent retrograde transport of BDNF-TrkB signalling endosomes to the soma is interrupted by local pausing at presynaptic boutons to modulate presynaptic plasticity (Andres-Alonso et al., 2019). This temporary stop is controlled by PKA-mediated phosphorylation of the Snapin cargo adaptor, which in turn releases the dynein motor complex to arrest the TrkB-BDNF signalling endosome. It is conceivable that molecular events at the presynaptic bouton might also trigger the activation of the SIPA1L2 PDZ domain. The activated SIPA1L2 PDZ domain could then bind Snapin to support its association with the BDNF-TrkB endosome after dynein motor complex release. Ultimately, the BDNF-TrkB signaling endosomes have to resume trafficking to the soma to trigger neuronal survival. After dephosphorylation and conformational changes switching the SIPA1L2 PDZ domain back to its inhibited state, Snapin would be able to again recruit the dynein motor complex to continue retrograde TrkB-BDNF signalling endosome trafficking.

## Supporting information

Supplementary text S1, supplementary figues S1-S5, supplementary table S1

## Acknowledgements

We acknowledge SOLEIL and the European Synchrotron Radiation Facility (ESRF) for the provision of synchrotron radiation facilities, and we would like to thank Andrew Thompson for assistance in using beamline PROXIMA 1 under proposal 20201046, and Daniele de Sanctis for assistance and support in using beamline ID30B under proposal number mx2461. ChatGPT (OpenAI; GPT-6) and Claude (Anthropic; Opus 5.5) were used between 07/2026 and 11/2026 to improve the grammar and phrasing of the manuscript, to write Python scripts for the structural and statistical analyses described in Section 2.5, and to write PyMOL 3 scripts with which the structure figures were rendered. All scripts were reviewed, executed, and checked by the authors, and all numerical results and figures were verified against the underlying data. The authors take full responsibility for the content of this article.

## Data availability

Coordinates and structure factors for the SIPA1L2 PDZ domain in the noCa, Li, and Ca forms have been deposited at the Protein Data Bank with PDB accession codes 35IQ, 35IS, and 35IT. The SIPA1L2 PDZ native holdup mass spectrometry proteomics data have been deposited in the ProteomeXchange Consortium via the PRIDE (Perez-Riverol et al., 2025) partner repository with the dataset identifier PXD086561.

