## Supplementary text S1, supplementary figues S1-S5, supplementary table S1 for "The crystal structure of the human SIPA1L2 PDZ domain reveals an atypically short, metazoan-conserved β2–β3 loop"

### Section S1:

Ordered metal ions decorate the ANXA2 chaperone in both the Ca and the Li form; no metal site is associated with the SIPA1L2 PDZ domain itself (Fig. S1). Six $Ca^{2+}$ ions are modeled in the Ca form ($Ca^{2+}$-1 - $Ca^{2+}$-6). Four of them are well defined ($Ca^{2+}$-1, $Ca^{2+}$-2, $Ca^{2+}$-3 and $Ca^{2+}$-6): evaluated in the full lattice, including symmetry-related contacts, they are five- to seven-coordinate, with Ca–O distances of 2.03–2.68 Å and bond-valence sums of 1.62–2.44 valence units, consistent with the nominal +2 (Zheng *et al.*, 2017). Three of these ($Ca^{2+}$-2, $Ca^{2+}$-3 and $Ca^{2+}$-6) are built on the canonical annexin architecture, in which three main-chain carbonyl oxygens of a glycine-rich loop are completed by a bidentate carboxylate donated by an acidic residue forty positions downstream. The remaining two ($Ca^{2+}$-4 and $Ca^{2+}$-5) are secondary surface sites that do not follow this architecture: each is coordinated by only one or two main-chain carbonyls together with an acidic side chain two or three residues away, and each presents an open, solvent-facing coordination sphere with no symmetry-related contacts. Their bond-valence sums (0.73 and 0.93 v.u.) and displacement parameters (50 and 53 $Å^2$, against a model mean of 34 $Å^2$) are those expected of partially occupied sites whose remaining coordination positions are completed by solvent not resolved at this resolution. Both are retained because omitting them degrades the refinement.

In the Li form, two of these same sites are occupied instead by $Li^{+}$ ($Li^{+}$-1 and $Li^{+}$-2) from the tri-lithium citrate mother liquor. Both lithium ions are coordinated exclusively by protein oxygen atoms and show markedly shorter metal–oxygen distances than their calcium counterparts, incompatible with an ordered water molecule at either site. $Li^{+}$-1, on the Gly1209/Arg1212/Gly1214–Glu1254 loop, is cleanly four-coordinate (1.89–1.98 Å; a second Glu1254 carboxylate oxygen lies at 3.04 Å, non-coordinating), with a bond-valence sum of 1.09 v.u. and a low displacement parameter (20 Å$^2$). $Li^+$-2, on the Met1285/Gly1287/Gly1289–Asp1329 loop, retains a monovalent-ion-like bond-valence sum (0.99 v.u.) but shows a more irregular, 4+1 coordination geometry: three tight donors at 1.71–2.24 Å and the first Asp1329 carboxylate oxygen at 2.07 Å, with the second Asp1329 oxygen at 2.82 Å, a longer, more marginal contact than the equivalent position at Li1. The single 1.71 Å contact (Gly1287 O) is shorter than typical for Li–O and disproportionately raises the site's bond-valence sum, so $Li^+$-2 is the less well-ordered of the two lithium sites; this pattern is consistent with a small, monovalent ion occupying a coordination geometry built for the larger, divalent $Ca^{2+}$ that ANXA2 normally binds at this position. Modeling either site as $Ca^{2+}$ or as $Na^+$ would give bond-valence sums of 3.8–4.2 v.u. and 2.5–2.7 v.u., respectively — grossly over-valent — so both assignments can be excluded at both sites. The two $Li^+$ sites occupy loops that host $Ca^{2+}$ in the Ca form; this coincidence provides an internal cross-validation of both assignments. Notably, $Li^+$ occupied them even though the protein buffer contained 20 mM $CaCl_2$. These ions belong to the crystallization chaperone and are incidental to SIPA1L2; they are reported as features of the crystallized fusion rather than as functional metal sites of the PDZ domain.

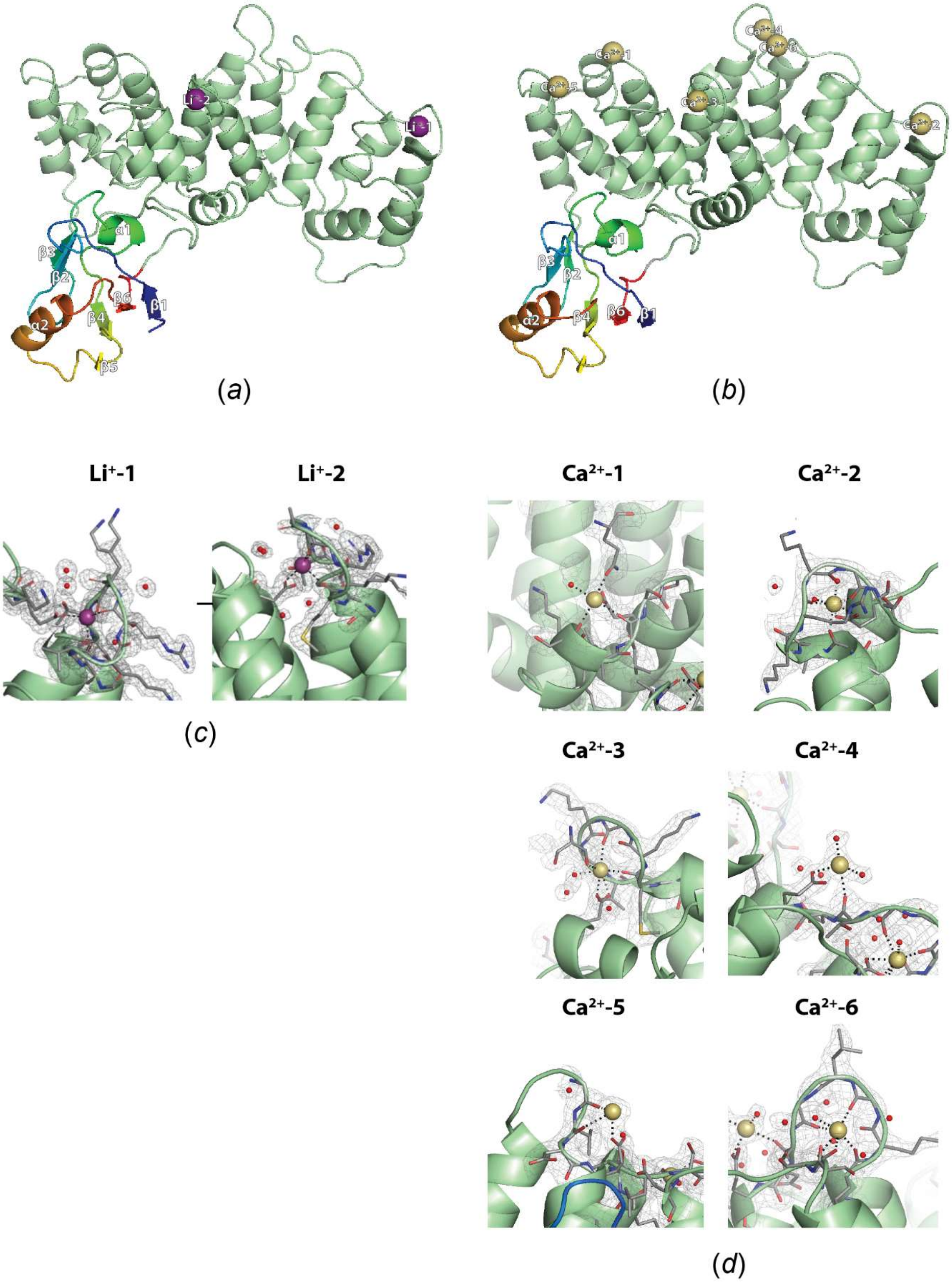
Li+-2
Li+-1
Ca2+-1
Ca2+-2
Ca2+-3
Ca2+-4
Ca2+-5
Ca2+-6
α1
α2
β1
β2
β3
β4
β5
β6
(a)
(b)
Li+-1
Li+-2
(c)
Ca2+-1
Ca2+-2
Ca2+-3
Ca2+-4
Ca2+-5
Ca2+-6
(d)

**Figure S1. Metal ion binding to the annexin crystallization tag. (*a*)** Li form ($P2_12_12_1$). (***b***) Ca form (C2). SIPA1L2 PDZ colored N→C-terminus and ANXA2 chaperone in palegreen. $Li^+$ ion are shown as purple spheres and $Ca^{2+}$ ions as olive-colored spheres. **(*c*)** Close-up of the two $Li^+$ binding sites. **(*d*)** Close-up of the six $Ca^{2+}$ binding sites. Water molecules are shown as red spheres. Coordination bonds are shown as black dotted lines. 2mFo−DFc electron density is contoured at 1.0σ and carved within 1.6 Å of the displayed atoms. For more detail, see Section S1.

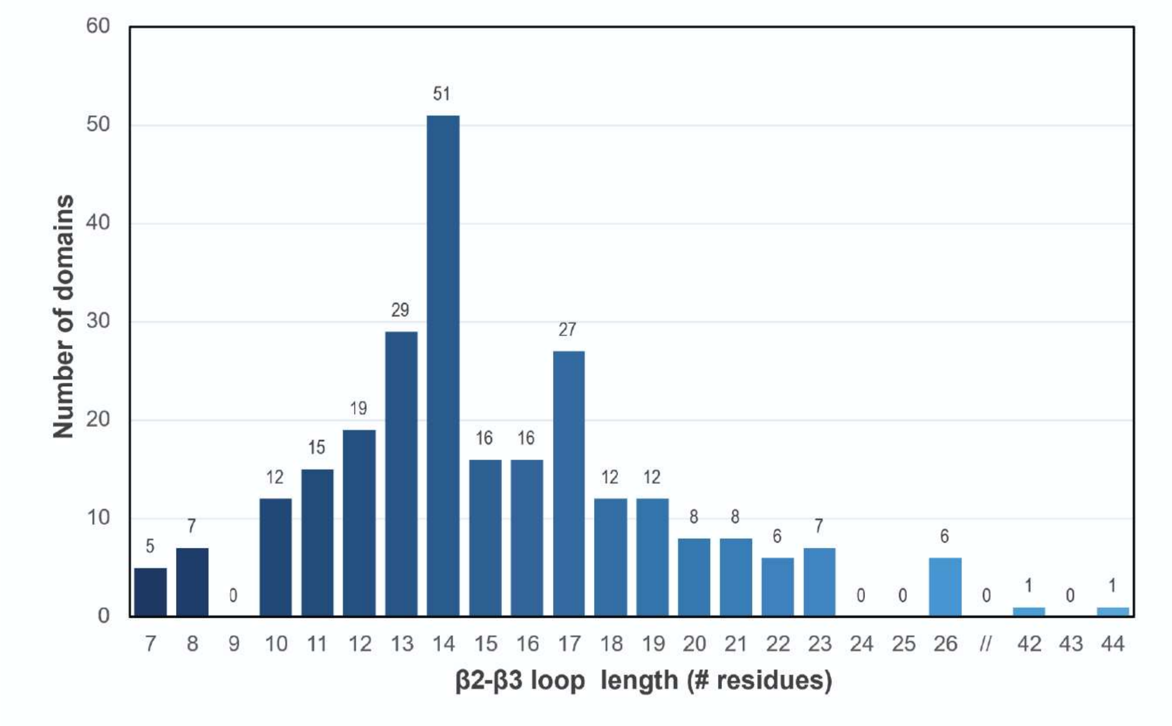

**Figure S2. β2–β3 loop-length distribution across the human PDZome.**

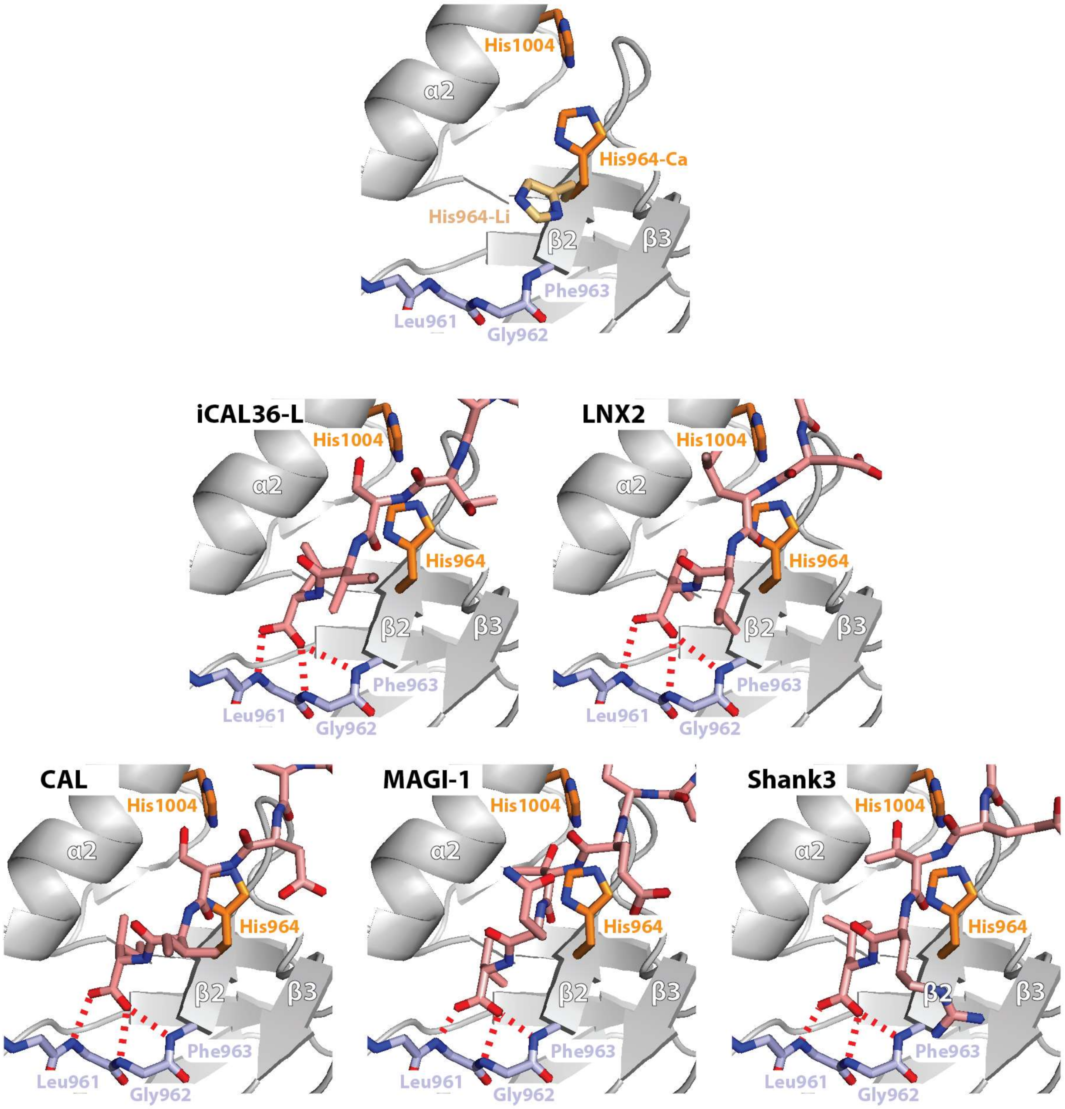

**Figure S3. His964 also obstructs the PBM binding groove in the SIPA1L2 PDZ Ca crystal form.** Top panel: Comparing the PBM binding groove of the Li to the Ca crystal form reveals a different side-chain conformation for His964. The His964 $\chi_1$ angle is −60° in the Li crystal form (pale orange), but −174° in the Ca crystal form (orange). Middle and lower panels: Docking of PBMs (salmon) of iCAL36-L (PDB-ID: 4e3b), LNX2 (PDB-ID: 7qct), CAL (PDB-ID: 4k72), MAGI-1 (PDB-ID: 2kpl) and Shank3 (PDB-ID: 5ovc) onto the SIPA1L2 PBM binding groove of the Ca crystal form. The SIPA1L2 PDZ domain (Ca form) is shown in grey cartoon representation. The main-chain of the carboxylate-binding loop (Leu961, Gly962, Phe963) is shown as light blue sticks. The α2 class-I reader His1004 and His964 are shown as orange sticks. The α2 helix and the β2 and β3 strands are labelled. In all cases, contacts with the canonical PBM binding motifs (carboxylate binding loop, α2 class-I reader) would be possible. Potential hydrogen bonds are shown as dashed red lines. Also in the altered side-chain conformation, His964 overlaps with the PBM main chain.

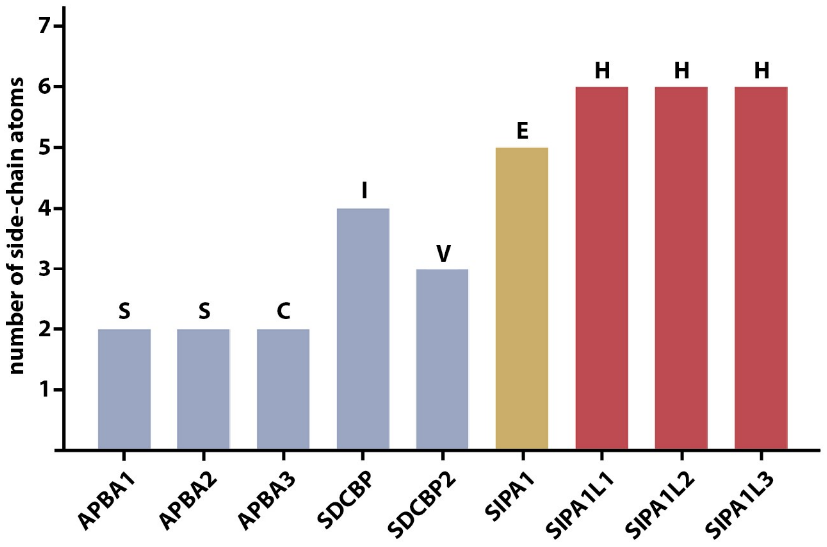

**Figure S4. The residue at the His964-equivalent position across human short-loop PDZ domains.** Bar height is the number of side-chain atoms (except hydrogens) of the residue structurally equivalent to SIPA1L2 His964 (one-letter code). A histidine occurs only in the SIPA1L subfamily (SIPA1L1/2/3, red). The PBM-binding short-loop domains carry a small residue (Ser in APBA1 and APBA2, Cys in APBA3, Ile in syntenin-1/SDCBP1, Val in SDCBP2, blue), and the divergent paralog SIPA1 (yellow) carries glutamate. The presence of smaller side chains (Ser, Cys, Ile, Val) correlates with PBM binding. The presence of larger side chains (Glu, His) correlates with the absence of identified PBM interactors (SIPA1, SIPA1L1/2/3).

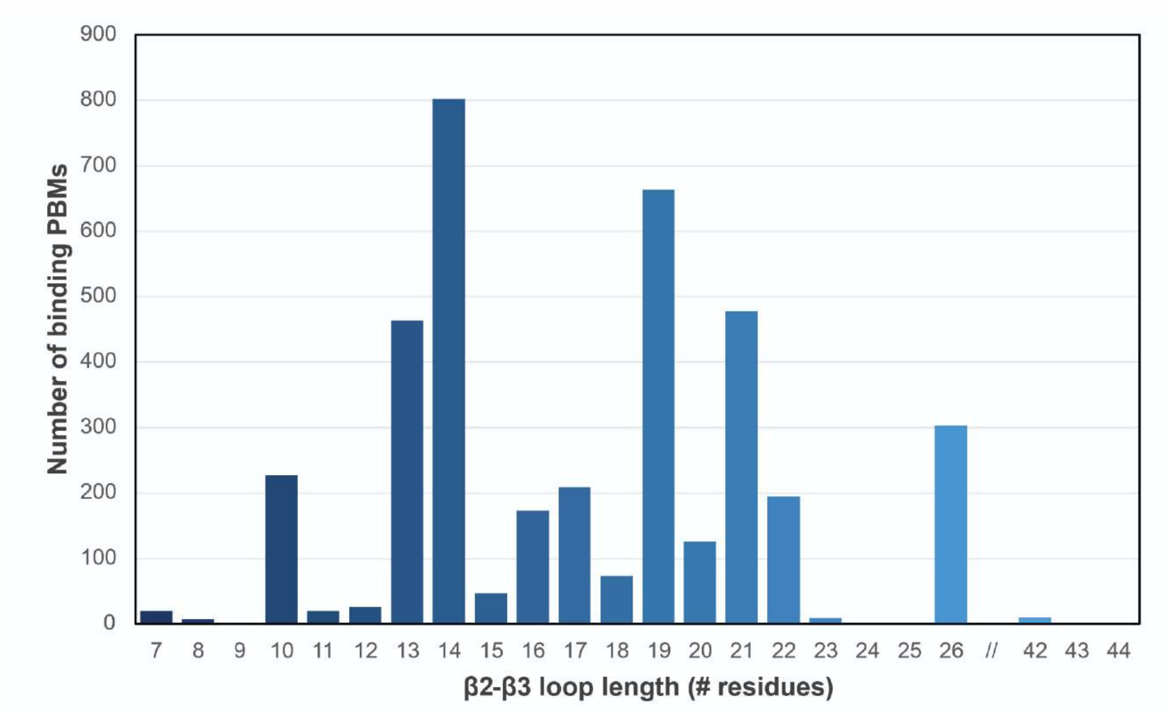

Number of binding PBMs
β2-β3 loop length (# residues)
0
100
200
300
400
500
600
700
800
900
//
42
43
44

(*a*)

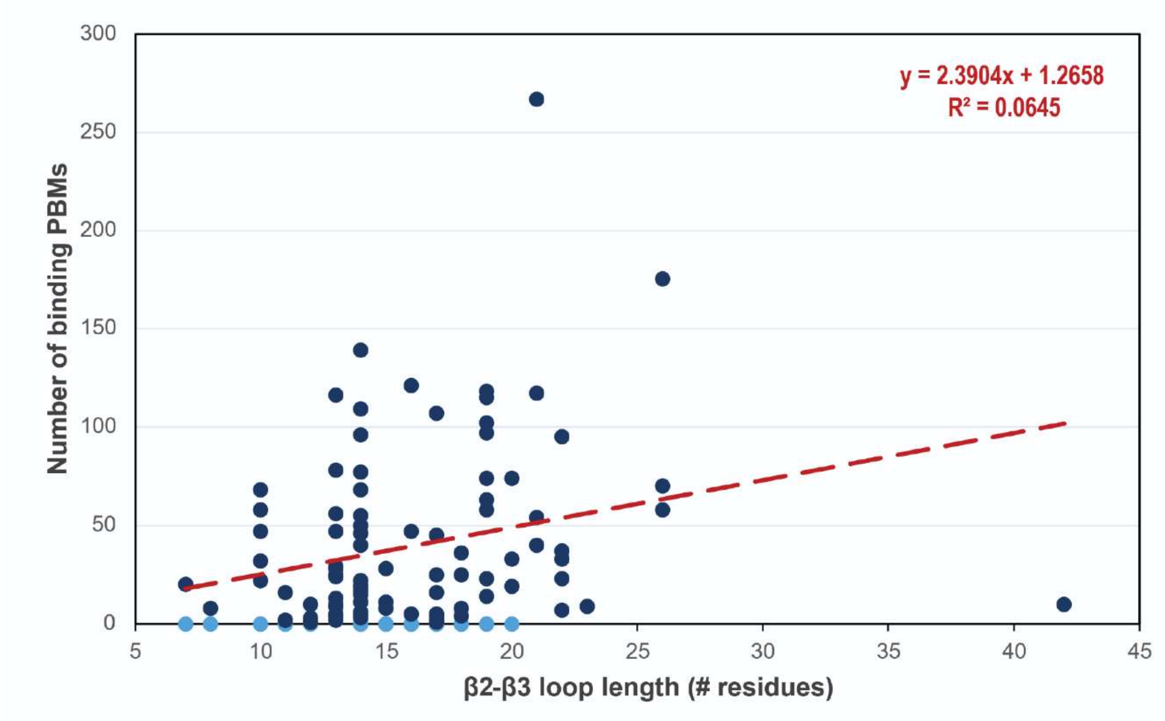

y = 2.3904x + 1.2658
R² = 0.0645
Number of binding PBMs
β2-β3 loop length (# residues)
0
50
100
150
200
250
300
5
10
15
20
25
30
35
40
45

(*b*)

**Figure S5. β2–β3 loop length versus reported PBM binding across the human PDZome**. **A)** Summed interactions per loop-length class and **B)** per-domain binder counts with linear fit. There is no correlation between the β2–β3 loop length and number of PBM interactors.

**Supplementary Table S1.** Metazoan homologs: species, PDZ sequence identity, Cα RMSD, β2–β3-loop sequence.

| Taxonomic group | Species | Protein Name | Sequence Identity of PDZ domain | RMSD of predicted PDZ | β2–β3 loop |
|---|---|---|---|---|---|
| **Mammal** | *Homo sapiens* | SIPA1L2 | 100 | 0.6 | FHVNFEGI |
| **Mammal** | *Pan troglodytes* | SIPA1L2 | 100 | 0.6 | FHVNFEGI |
| **Mammal** | *Mus musculus* | SIPA1L2 | 100 | 0.6 | FHVNFEGI |
| **Monotremata** | *Ornithorhynchus anatinus* | SIPA1L2 | 97 | 0.6 | FHVNFEGI |
| **Amphibia** | *Leptobrachium leishanense* | SIPA1L2 | 96 | 0.6 | FHVNFEGI |
| **Teleostei** | *Megalops atlanticus* | SIPA1L2 | 96 | 0.6 | FHVNFEGV |
| **Jawless Fish** | *Eptatretus burgeri* | *SIPA1L1** | 88 | 0.6 | FHVNYEGV |
| **Basal chordates** | *Branchiostoma floridae* | SIPA1L2 | 85 | 0.8 | FHVDLEGM |
| **Echinoderms** | *Strongylocentrotus purpuratus* | SIPA1L2 | 75 | 0.5 | FHVHYQGI |
| **Arthropoda** | *Amphibalanus amphitrite* | SIPA1L2 | 61 | 0.7 | FHVQSDGL |
| **Nematoda** | *Caenorhabditis elegans* | *SIPA1** | 43 | 0.8 | FHVQDEGV |
| **Nematoda** | *Plectus sambesii* | SIPA1L2 | 55 | 0.7 | FHLHEEGV |
| **Annelida** | *Ridgeia piscesae* | SIPA1L2 | 79 | 0.7 | FHIHYDGI |
| **Mollusca** | *Elysia crispata* | SIPA1L2 | 72 | 0.6 | FHIQSEGI |
| **Cnidaria** | *Stylophora pistillata* | SIPA1L2 | 68 | 0.7 | FHVQFEGL |
| **Placozoa** | *Trichoplax sp. H2* | *SIPA1L1** | 25 | 0.8 | LHIYPDGI |
| **Porifera** | *Geodia barretti* | *SIPA1L1** | 27 | 0.7 | FSINEVCE |
| **Mammal** | *Homo sapiens* | *SIPA1L1** | 92 | 0.6 | FHVNYEGI |
| **Mammal** | *Homo sapiens* | *SIPA1L3** | 84 | 0.7 | FHVKYDGT |
| **Mammal** | *Homo sapiens* | *SIPA1** | 43 | 0.8 | FEVDAEGF |
